# Lsr2 is a nucleoid-associated protein that exerts pleiotropic effects on mycobacterial cellular processes

**DOI:** 10.1101/2020.04.27.063487

**Authors:** Marta Kołodziej, Damian Trojanowski, Katarzyna Bury, Joanna Hołówka, Mariola Paściak, Weronika Matysik, Hanna Kąkolewska, Helge Feddersen, Giacomo Giacomelli, Marc Bramkamp, Igor Konieczny, Jolanta Zakrzewska-Czerwińska

**Affiliations:** Department of Molecular Microbiology, Faculty of Biotechnology, University of Wrocław, Wrocław, Poland; Department of Molecular and Cellular Biology, Intercollegiate Faculty of Biotechnology, University of Gdansk and Medical University of Gdansk, Gdansk, Poland; Department of Immunology of Infectious Diseases, Hirszfeld Institute of Immunology and Experimental Therapy, Polish Academy of Sciences, Weigla 12, 53-114, Wroclaw, Poland; Christian-Albrechts-Universität zu Kiel, Institut für allgemeine Mikrobiologie, 24118 Kiel, Germany

## Abstract

Lsr2 is involved in maintaining chromosome structure in asymmetrically dividing mycobacteria and is essential in the tubercle bacillus (*M. tuberculosis*) during infection. Here, we report that a lack of Lsr2 profoundly impacts the mycobacterial cell morphology and the properties of the cell envelope resulting in the formation of smooth, short and antibiotics sensitive cells. Lsr2 forms large and dynamic nucleoprotein complexes *in vivo* and deletion of *lsr2* gene exerts a profound effect on the replication time and replisome dynamics. We suggest that the Lsr2 nucleoprotein complexes may contribute to maintaining the proper organization of the newly synthesized DNA. Moreover, we demonstrate that the N-terminal oligomerization domain of Lsr2 is indispensable for the formation of nucleoprotein complexes *in vivo*. Collectively, our results indicate that Lsr2 exerts a pleiotropic effect on cellular processes and appears to be an attractive target for the development of a novel antitubercular drugs.

## Introduction

To fit chromosomes into the tiny volume of a prokaryotic cell (bacteria and Archaea) or a eukaryotic nucleus, the DNA of each chromosome must be efficiently compacted and organized (Fisher et al., 2013; Le et al., 2013; Toro & Shapiro, 2010; X. Wang, 2006). Although chromosomal DNA is condensed by different factors, including DNA supercoiling (Hsu, Chung, & Li, 2006; Kahramanoglou et al., 2011; Thanbichler, Wang, & Shapiro, 2005), it is primarily organized by basic DNA-associated proteins such as histones (in eukaryotes and Archaea) or nucleoid-associated proteins (NAPs; in bacteria) (Luijsterburg et al., 2008; Nolivos et al., 2016; Prieto et al., 2012a; Tolstorukov et al., 2016). In addition to their substantial impact on the chromosomal architecture, histones and NAPs may also act as global transcription factors (Badrinarayanan, Le, & Laub, 2015; Berger et al., 2010; Kahramanoglou et al., 2011; Luijsterburg et al., 2006; K. Singh, Milstein, & Navarre, 2016).

Bacteria possess diverse sets of NAPs that compact chromosomes by bending, bridging, or wrapping the DNA (Beloin et al., 2003; Dame, Noom, & Wuite, 2006; Stavans & Oppenheim, 2006). Their interactions with DNA are highly dynamic; they depend on the growth phase and environmental cues and adjust global gene expression to different conditions (Azam et al., Iwata, 1999; Karas, Westerlaken, & Meyer, 2015; Moore et al., 2015; Prieto et al., 2012b; Ushijima, Ohniwa, & Morikawa, 2017). H-NS (histone-like nucleoid-structuring protein), which is one of the best characterized NAPs, plays important roles in chromosomal DNA organization and gene silencing (Dame, 2005; Dillon & Dorman, 2010; Wang et al., 2011). H-NS possesses two functional domains, an N-terminal dimerization domain and a C-terminal DNA binding domain, that are separated by a flexible linker (Gordon et al., 2011). Two distinct DNA-binding modes have been reported for H-NS: i) DNA bridging by intra- and intermolecular DNA binding, and ii) DNA stiffening by polymerization along the DNA double helix to form a rigid filament (Dame et al., 2006). Both activities can silence gene expression by limiting the access of RNA polymerase, while DNA bridging also facilitates chromosome compaction (Lim et al., 2012; Maurer, Fritz, & Muskhelishvili, 2009; Navarre et al., 2007).

The *Mycobacterium* genus encompasses not only human pathogens that have enormous impact on global health (i.e., *Mycobacterium tuberculosis, M. leprae*), but also the saprophytic species, such as *Mycobacterium smegmatis,* which is a model organism for studies on the cell biology of tubercle bacilli. Mycobacteria possess a unique multilayered cell envelope and, unlike other rod-shaped bacteria, incorporate peptidoglycan precursors at their cell tips (Abrahams & Besra, 2018; García-Heredia et al., 2018; B. Singh et al., 2013). The daughter cells that inherit the old cell pole (with the preassembled elongation machinery) elongate faster than their siblings that inherit the new pole (Aldridge et al., 2012; Baranowski et al., 2018; Logsdon & Aldridge, 2018). Moreover, mycobacteria often divide asymmetrically, generating unequally sized daughters (Kieser & Rubin, 2014). Unlike the situation in other extensively studied bacteria, such as *Escherichia coli*, the chromosome is positioned asymmetrically within the mycobacterial cell throughout the entire cell cycle, and its structure is maintained by a unique set of NAPs (Kriel et al., 2018). Due to their relative lack of sequence homology to their *E. coli* counterparts, most of the mycobacterial NAPs have only recently begun to be identified. Lsr2, one of the principal mycobacterial NAPs, is believed to function similarly to H-NS: It is a structural homolog of H-NS, preferentially binds AT-rich sequences, and is able to bridge distant DNA fragments or form a rigid nucleoprotein filament *in vitro* (Chen et al., 2008; B. R. G. Gordon et al., 2011; Blair R G Gordon, Imperial, Wang, Navarre, & Liu, 2008; Qu, Lim, Whang, Liu, & Yan, 2013; Summers et al., 2012).

To date, most of the *in vivo* studies on Lsr2 have examined its influence on gene expression (Gordon et al., 2011). In *M. tuberculosis*, Lsr2 regulates (mainly represses) the transcription of multiple genes involved in a broad range of cellular processes, including cell wall synthesis, virulence, and the adaptation to changing oxygen levels (Bartek et al., 2011; Roberto Colangeli et al., 2007). In the present study, we sought to explore the biological role of Lsr2 at the single-cell level. We demonstrate that deletion of the *lsr2* gene has profound impacts on cell morphology and the properties of the cell envelope, including lipid content, resulting in the formation of smooth and short cells. Time lapse-fluorescent microscopy (TLFM) experiments revealed that Lsr2 forms a highly dynamic nucleoprotein complex(es) and influences both the dynamics of replication machinery and the duration of DNA synthesis. Finally, using TLFM and photoactivated localization microscopy (PALM), we show that the N-terminus is indispensable for the formation of the Lsr2-DNA complex.

## Results

### Lack of Lsr2 affects cell morphology

The mycobacterial Lsr2 protein has been extensively studied in the last decade, but this work has mainly involved *in vitro* approaches, with the *in vivo* studies remaining limited to bulk-culture observations of the bacterial population. Therefore, we decided to examine the influence of Lsr2 on cellular processes at the single-cell level. We constructed an *M. smegmatis* strain with deletion of the *lsr2* gene (*msmeg_6092*). Consistent with previous reports on similar strains (Arora et al., 2008; Chen et al., 2006; Gordon et al., 2008), the *M. smegmatis* Δ*lsr2* strain (Δ*lsr2*) exhibited changes in colony morphology, biofilm formation, and sliding motility in comparison to the wild-type mc^2^ 155 strain (WT). The strain lacking Lsr2 formed round and smooth colonies on 7H10 or NB agar plates, was unable to form biofilm, and could spread on the surface of solid medium (for details see Fig. S1).

To analyze cell morphology, we stained Δ*lsr2* and WT cells using either a lipophilic membrane dye (FM 5-95), fluorescent D-alanine (NADA) and/or fluorescent trehalose (TMR-Tre; for details see Materials and Methods). We found that the cells with *lsr2* deletion were about 42% shorter (2.9 +/− 0.6 vs. 5.0 +/− 1.4 μm, respectively, n = 100, *p* = 2×10^−16^) and 30% wider (0.94 +/− 0.9 μm vs. 0.71 +/− 0.07 μm, respectively, n = 80, *p* = 2×10^−16^) than the WT cells (mean +/− standard deviation, n – observation number, *p* value by pairwise t test with the pooled SD) (Fig. 1A).

**Fig. 1.**
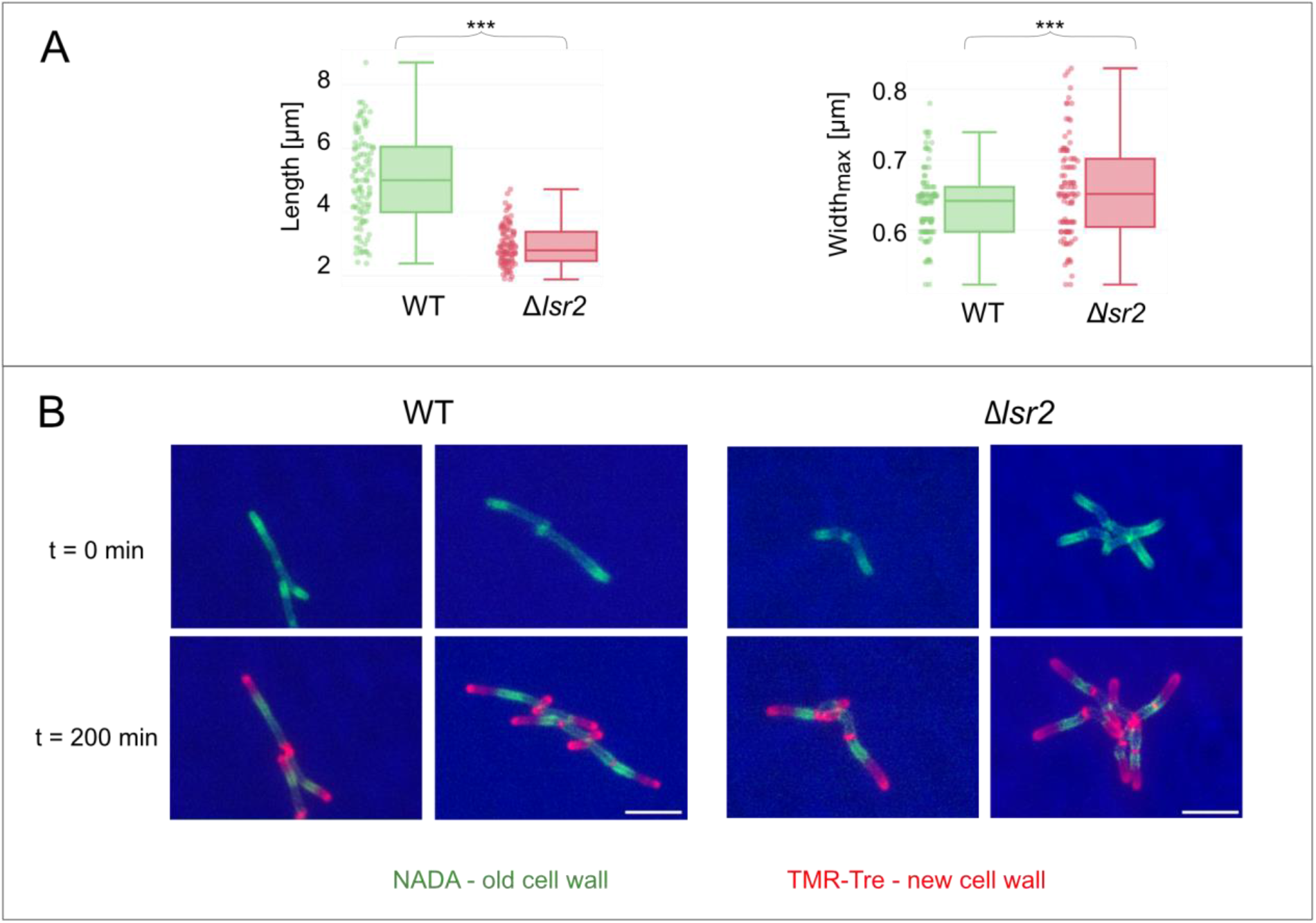
Influence of lsr2 deletion on mycobacterial cell morphology. **(A)** Comparison of the cell length (left) and width (right) between the wild-type *M. smegmatis* mc^2^ 155 (WT) and Δ*lsr2* strains (n = 100). Statistical significance was defined as \*\*\**p* < 0.0005 (parametric double-sided t-test with pooled SD). **(B)** Micrographs showing representative cells of the WT and Δ*lsr2* labeled with NADA and TMR-Tre dyes. Bar, 5 μm.

To further investigate the cell morphology of Δ*lsr2* mutant cells, we performed atomic force microscopy (AFM) analysis of the cell stiffness and surface roughness. In general, our AFM observations (Fig. 2A) confirmed that Δ*lsr2* cells were shorter and wider than WT cells. To estimate the stiffness, cells were immobilized on a PDMS slide and imaged in the PeakForce QNM mode. We then determined Young’s modulus, which defines the relationship between the mechanical tension applied on a material and the resulting strain, and is measured in Pascals (Pa, N m^−2^). The Young’s modulus calculated for Δ*lsr2* cells was higher than the corresponding value for WT cells (2.6 +/− 0.6 MPa vs. 1.1 +/− 0.2 MPa, respectively, n = 30, *p* < 2×10^−16^)(Fig. 2B, top panel), indicating that the mutant cells are more rigid (presumably due to a high turgor pressure, see Discussion). Further investigation of cell morphology using AFM imaging was performed using the PeakForce Tapping mode. This analysis showed that the cell surface of the Δ*lsr2* strain was smoother than that of the WT strain; the calculated Ra parameter (arithmetic mean deviation of the roughness profile) was 3.33 +/− 1.81 and 4.68 +/−1.25 for Δ*lsr2* and WT cells, respectively (Fig. 2B, bottom panel; 50 cells were analyzed for each strain).

**Fig. 2.**
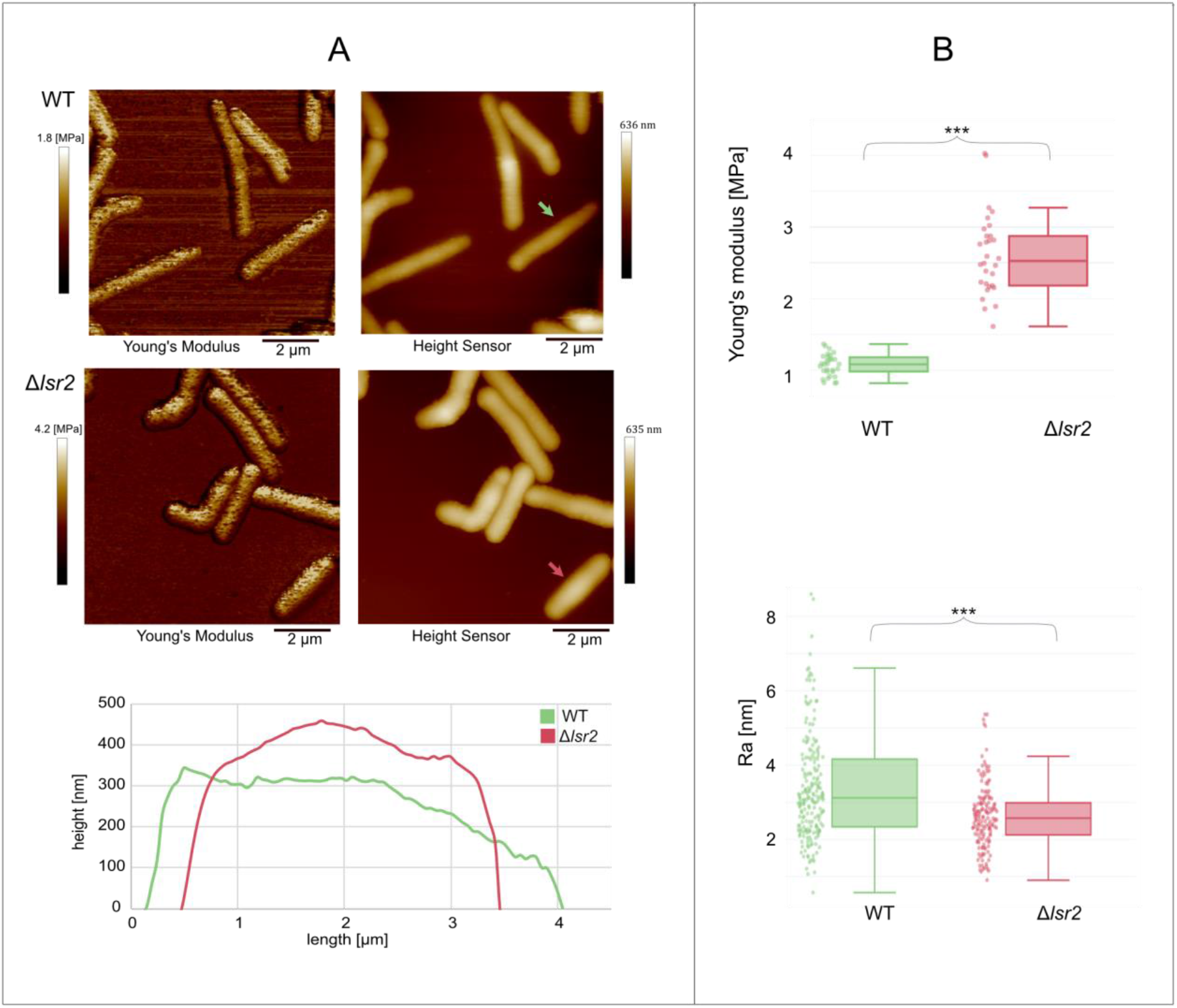
AFM analysis of WT and Δ *lsr2 M. smegmatis* cells. **(A)** Young’s modulus and height sensor images of mc^2^ 155 and Δ*lsr2* cells (top panel). The height profiles of representative cells of each strain (bottom panel) are indicated by the green and red arrows, respectively. **(B)** Young’s modulus (a measure of elasticity) and Ra parameter (arithmetic mean deviation of the roughness profile) were compared for WT and Δ*lsr2* cells. To measure Young’s modulus, five uniformly spaced points were selected on each bacterial cell (n = 30, top panel). Ra was calculated using measurements taken from three or four 500-nm lines per cell (n = 50, bottom panel; statistical significance was defined as \*\*\**p* < 0.0005; parametric double-sided t-test with pooled SD).

**Fig. 3.**
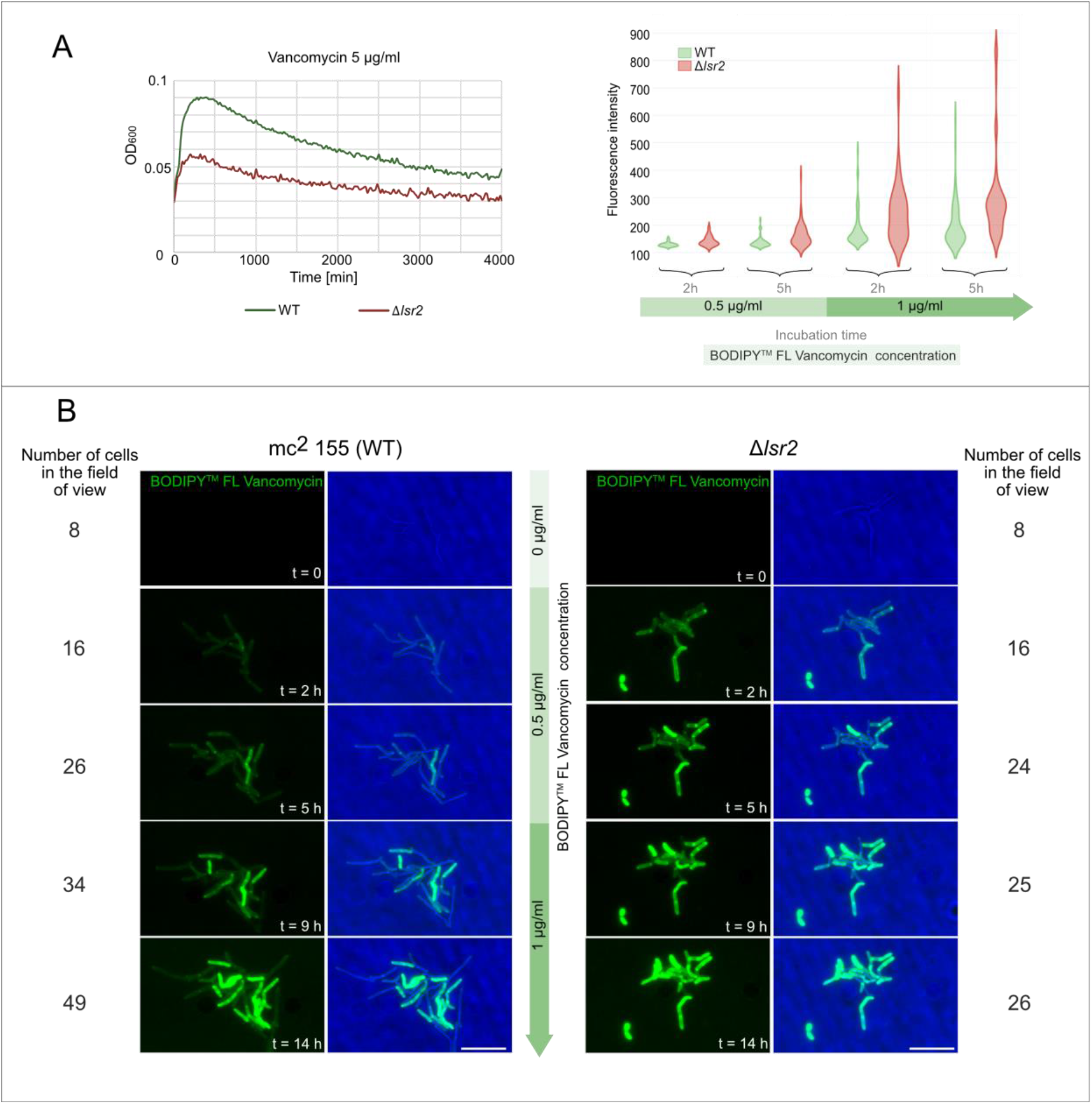
Comparison of vancomycin susceptibility and incorporation of Vancomycin-BODIPY into the cell wall of WT versus Δlsr2 cells. **(A)** Growth curves of the *M. smegmatis* strains in the presence of 5 μg/ml vancomycin (left panel). The fluorescence intensity of Vancomycin-BODIPY incorporated into the cell wall was measured at four different time points (right panel). Maximal fluorescence intensity was observed for the lateral cell wall away from the division site (n = 50 to 67 cells for each time point). **(B)** TLMM (time-lapse microfluidic microscopy) analysis of Vancomycin-BODIPY incorporation. Cells were exposed to 0.5 μg/ml Vancomycin-BODIPY for 7 h, whereupon the concentration was increased to 1 μg/ml. The number of surviving cells is shown on the side of the micrographs. Bar, 5 μm.

The observed differences in the cell surface roughness may reflect changes in the properties of the cell envelope particularly in the composition of the outer leaflet of the mycomembrane. Indeed, the pathogenic nontuberculous species (*M. abscessus* or *M. kansasii*) depending on the level of surface-associated glicolipids (glycopeptidolipid (GPL) or lipooligosaccharides (LOS)) exhibit either a smooth (S) or a rough colony morphotype (R) (Howard et al., 2006; Medjahed, Gaillard, & Reyrat, 2010; Roux et al., 2016). Various hypotheses regarding the influence of Lsr2 on mycobacterial cell wall composition have been proposed, but all of them assume that Lsr2 regulates genes encoding proteins involved in the synthesis of cell envelope components, including the outer leaflet that is heterogeneous among mycobacteria and consists of different lipids, lipoglycans and LOSs (Chiaradia et al., n.d.). Interestingly, microarray analysis of Δ*lsr2 M. smegmatis* strain (*lsr2* transposon mutant)(Colangeli et al., 2007) demonstrated that MSMEG_4727 gene encoding Mas-like Pks (mycocerosic acid synthase – like polyketide synthase) involved in the synthesis of LOS was expressed 6-fold higher than in the WT (mc^2^ 155) (Etienne et al., 2009). To validate the microarray results, we performed qRT-PCR using primers specific for the MSMEG_4727 gene (Table S3). Indeed, our results confirmed that the transcription level of MSMEG_4727 in the Δ*lsr2* strain compared to the wild type strain was 17-fold increased (Fig. 4A). Moreover, TLC analysis of methanol-soluble crude lipids (for details see Supplemental Experimental Procedures) revealed presence of the additional glycolipid spot in the Δ*lsr2* strain. This lipid compound was also observed in the Δ*lsr2* complemented strain, but its amount was significantly lower than in the Δ*lsr2* strain (Fig. 4B). To further confirm the LOSs presence in the Δ*lsr2* strain, we performed MALDI-TOF analysis of methanol-soluble crude lipids (Fig. S2C). Despite the presence of high-abundance GPLs in the wild type and Δ*lsr2*, two peaks (at m/z 1689.9 and 1429.9) corresponding to lipooligosacharides class (LOS A and LOS B1, respectively) (Etienne et al., 2005) were identified in the Δ*lsr2* strain (see Fig. S2C).

**Fig. 4.**
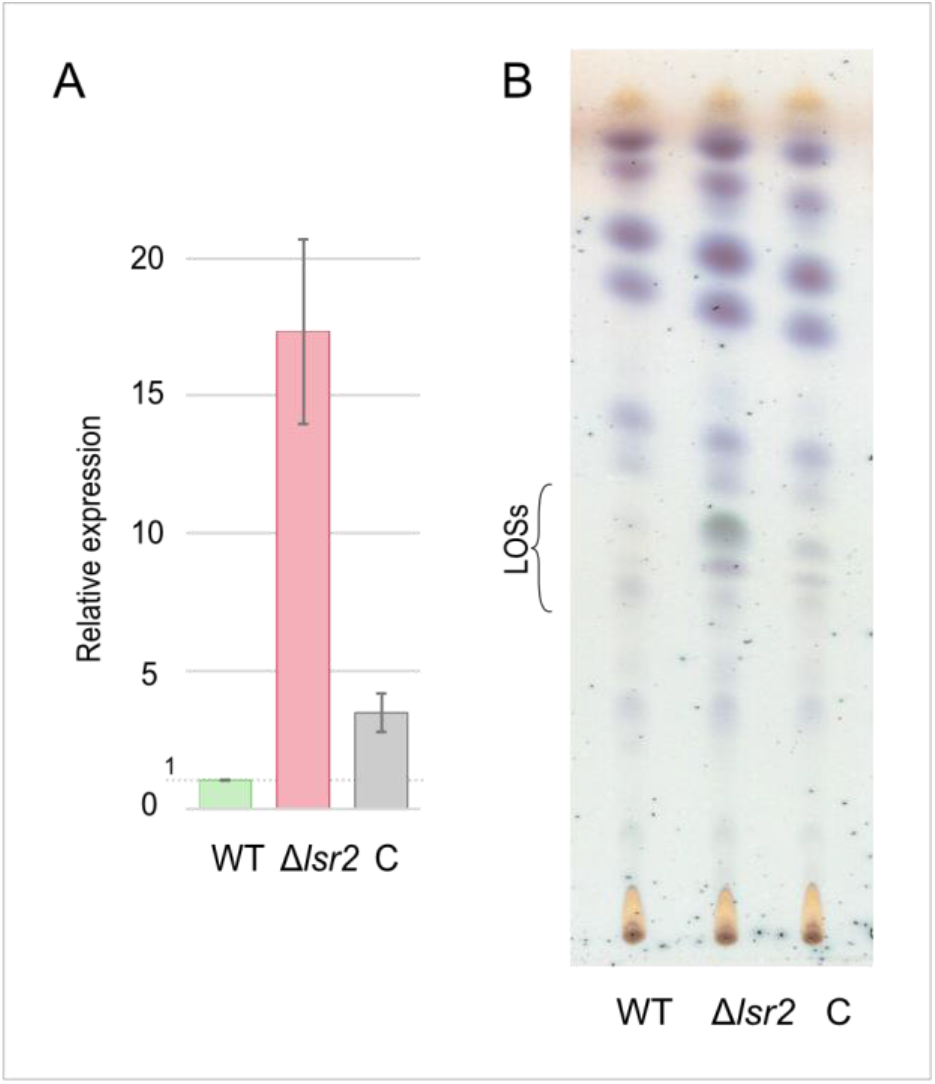
TLC analysis of LOSs plus RT-PCR of MSMEG_4727. (A) The relative transcription of MSMEG_4727 gene encoding mas-like PKS involved with LOSs synthesis, in *M. smegmatis* wild type mc^2^ 155 strain (WT), Δ*lsr2* and the complemented strain (C; Δ*lsr2*_pNAT*lsr2*), calculated using RT-qPCR analysis. **(B)** TLC analysis of crude methanol-soluble lipid fractions of *M. smegmatis* wild type mc^2^ 155 strain (WT), Δ*lsr2* and the complemented strain (C; Δ*lsr2*_pNAT*lsr2*). The 100μg of each sample were applied. Solvent system: chloroform-methanol (90:10, v/v). Detection: 0.2% anthrone in concentrated sulfuric acid.

We hypothesized that changes in the LOSs level, which were presumably mirrored by differences in the smoothness of Δ*lsr2* cells may have directly altered the antibiotics penetration. LOS is an amphipathic molecule that consists of a hydrophobic lipid part (that anchors the LOS to the outer membrane) and a hydrophilic carbohydrate part (Etienne et al., 2009). To test this hypothesis, we used green fluorescent-labeled vancomycin (Vancomycin-BODIPY; BODIPY^TM^FL Vancomycin), a hydrophilic antibiotic. Growth-curve analysis showed that the Δ*lsr2* strain was more susceptible to unlabeled vancomycin than the wild-type strain (Fig. 3A, left panel). Moreover, during TLFM experiments, we observed that Vancomycin-BODIPY was incorporated faster into the cell wall of the Δ*lsr2* strain. The fluorescent intensity measured at four different time points during incubation with Vancomycin-BODIPY was higher in the Δ*lsr2* strain than in the WT strain (Fig. 3A right panel, Fig. 3B).

Together, these findings indicate that deletion of the *lsr2* gene has a striking effect on single-cell morphology, triggering changes in cell size and cell envelope properties.

### Deletion of lsr2 results in a shortened C-period and altered replisome dynamics

Since the lack of Lsr2 protein was found to have a profound effect on cell envelope properties and cell morphology (i.e., triggering the formation of shorter cells), we sought to elucidate the impact of Lsr2 on the cell cycle. Using time-lapse microscopy, we compared the cell elongation rate (the increase of cell length per hour [μm × h-1] calculated by comparing the cell length measured shortly after cell division and at 50 min thereafter) and the duration of individual cell cycle periods (B, C, and D) in the Δ*lsr* and WT strains. Our results showed that the lack of Lsr2 resulted in a decreased cell elongation rate (Δ*lsr2* 1.1 +/− 0.3 μm×h^−1^ vs. WT 1.5 +/− 0.5 μm×h^−1^, n = 50, *p* = 1.7×10^−4^, Fig. 5A).

**Fig. 5.**
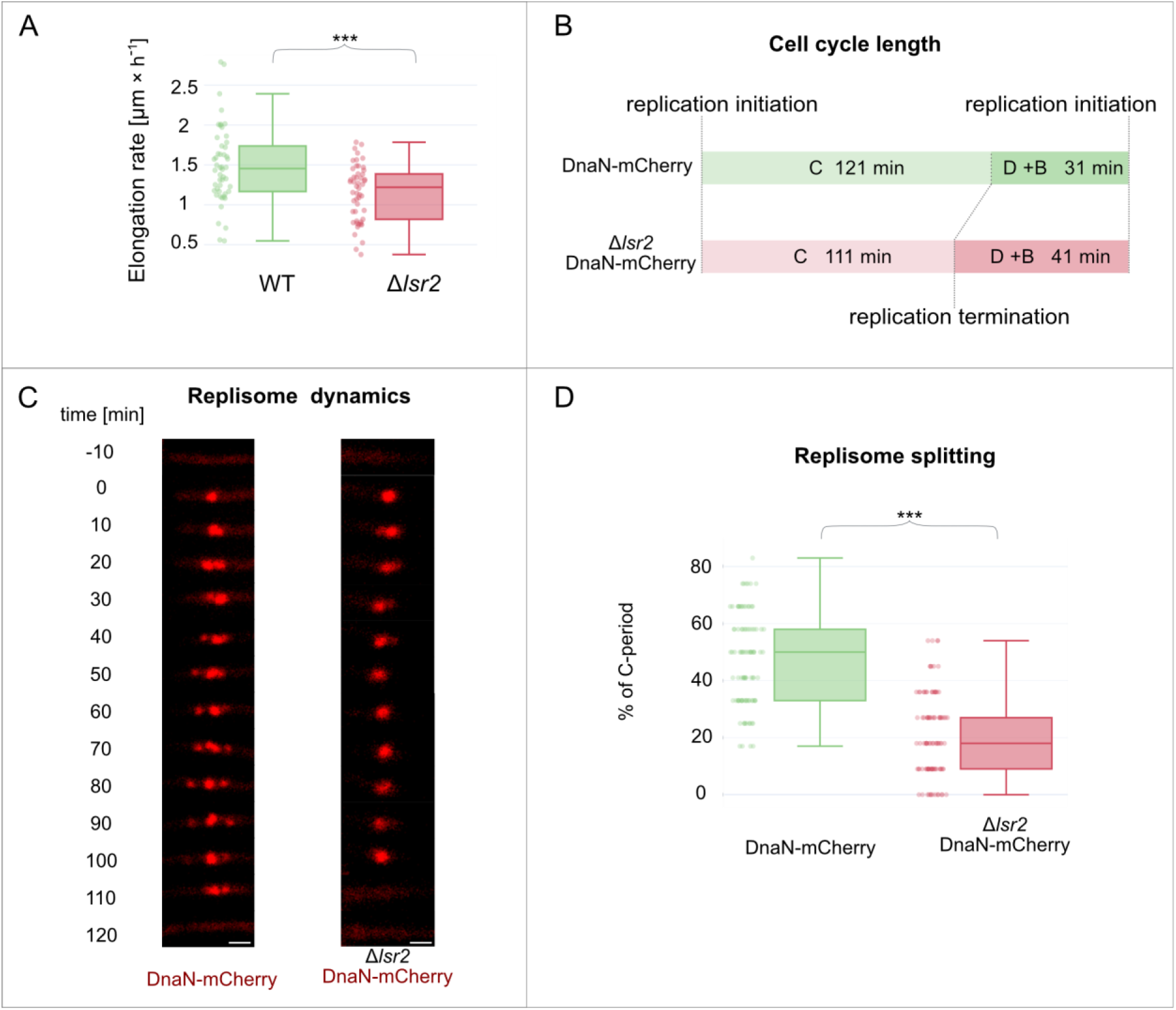
Influence of lsr2 deletion on the cell cycle and replisome dynamics. **(A)** Box-plot comparison of the elongation rate of WT versus Δ*lsr2* cells (n = 50, statistical significance was defined as \*\*\**p* < 0.0005, as assessed with a parametric double-sided t-test with pooled SD). **(B)** Schematic representation of the mycobacterial cell cycle in the WT and *lsr2* deletion strains. **(C)** Real-time microscopic analysis of replisome dynamics during one replication event. In WT cells, replisomes frequently divide and merge; in Δ*lsr2*, in contrast, the replisomes form one fluorescent focus for most of the C-period. Bar, 1 μm. **(D)** Replisome splitting expressed as the duration when the DnaN-mCherry marked replisomes were observed as more than one fluorescent focus, calculated as a percent of the total replication duration. (n = 100, \*\*\**p* < 0.0005; parametric double-sided t-test with pooled SD).

To examine the influence of Lsr2 on cell cycle parameters and the dynamics of chromosome replication (as a key process of the cell cycle), we constructed the Δ*lsr2*_DnaN-mCherry strain (for details see Table S1), which allowed us to perform real-time monitoring of DNA replication in the background of *lsr2* deletion. DnaN, a DNA polymerase III subunit (β-clamp), is frequently fused with fluorescence proteins (FP) to analyze the dynamics of the replisome (the multiprotein complex involved in DNA synthesis). The appearance and disappearance of DnaN-FP fluorescent foci correspond to the initiation and termination of replication, respectively, and the time between these events reflects the duration of replication (the C-period) (Santi & McKinney, 2015; Trojanowski et al., 2015). The constructed Δ*lsr2*_DnaN-mCherry strain exhibited a colony morphology and growth rate similar to those of the Δ*lsr2* strain (data not shown). TLFM analysis revealed that the duration of chromosome replication was shorter in the Δ*lsr2_*DnaN-mCherry strain than in the DnaN-mCherry control strain (111+/− 16 min and 121 +/− 13 min for Δ*lsr2_*DnaN-mCherry and DnaN-mCherry, respectively, n = 100, *p* = 3.7×10^−6^, Fig. 5B), whereas the time between replication termination and replication initiation in daughter cells (the D+B period) was longer (44 +/− 19 min and 31 +/− 11 min for Δ*lsr2_*DnaN-mCherry and DnaN-mCherry, respectively, n = 200, *p* = 6×10^−15^). Thus, the overall duration of the cell cycle (B+C+D) remained unchanged (Δ*lsr2*, 152 min vs. DnaN-mCherry control strain, 152 min; see Fig. 5B).

Since deletion of *lsr2* led to shortening of the C-period, we questioned how Lsr2 might influence replisome dynamics during the cell cycle. In *Mycobacterium*, replisomes appear to oscillate around each other, frequently merging and splitting (Trojanowski et al., 2015). This dynamic was disturbed in the Δ*lsr2_*DnaN-mCherry strain, as split replisomes were observed in these cells for only 21 +/− 14% of the replication time, compared to 47 +/− 16% in the DnaN-mCherry strain (n = 100, *p* < 1×10^−5^; Fig. 5CD).

We also analyzed the impact of Lsr2 on chromosome segregation during the cell cycle. For this purpose, we used Δ*lsr2*_ParB-mNeonGreen (for details see Table S1) and Δ*lsr2_*HupB-EGFP, which allowed us to monitor the segregation of the newly replicated *oriC* regions and chromosome separation, respectively. ParB binds in the vicinity of *oriC* and forms a nucleoprotein complex called the segrosome; thus, it can serve as an *oriC* localization marker (Ginda et al., 2013; Holówka et al., 2018). TLFM analysis of both strains showed that there was no difference in the *oriC* positioning or duplication time, the latter of which was defined as the period between two consecutive segrosome doublings (147 +/− 19 min for ParB-mNeonGreen vs. 147 +/− 16 min for Δ*lsr2_*ParB-mNeonGreen, n = 92, *p* = 1). The chromosome separation time was also unaffected (147 +/− 17 min for HupB-EGFP vs. 150 +/−19 min for Δ*lsr2_*HupB-EGFP, n = 100, *p* = 0.18). These data further confirmed our earlier finding that deletion of the *lsr2* gene does not alter the cell cycle duration.

### Lsr2 exhibits a highly dynamic localization during the cell cycle

Given that Lsr2 forms extensive oligomers upon DNA binding *in vitro* (Colangeli et al., 2007; Summers et al., 2012) and affects replisome dynamics (as shown immediately above), we decided to analyze its subcellular localization during the cell cycle. For this purpose, we used fluorescent-tagged Lsr2 protein. To avoid potential artifacts in our localization of subcellular Lsr2, we constructed and tested strains that produced Lsr2 fused with different fluorescent proteins (Lsr2-FP): EGFP, mCherry, mTurquoise2, and Dendra2 (for details see Table S1). TLFM revealed that in each analyzed strain, depending on the cell cycle stage, the Lsr2-FP was visible as one or two discrete and bright major foci accompanied by several unstable minor foci (Fig. 6 and data not shown). Additionally, we observed a dispersed fluorescence signal along the cell. The major Lsr2-FP focus (foci) was (were) highly dynamic, frequently changing in size, shape, and intensity (from more compact and dense to ragged and blurred; see Fig. 6A, Video S1). In the case of the Lsr2-EGFP fusion, however, the foci were more clustered and the duplication of the major focus was delayed compared to the other fusions (data not shown). Thus, our observations are consistent with the subcellular localization of H-NS (a structural homolog of Lsr2) fused with GFP. Although GFP (and also EGFP) exhibits very weak dimerization activity, it may act like a “velcro” (Wang et al., 2014), and thus could potentially assemble Lsr2-GFP (or H-NS-GFP) into larger complexes. Therefore, in further studies, we decided to use Lsr2 fused with other fluorescent proteins, such as mCherry, mTurquoise2, or Dendra2.

**Fig. 6.**
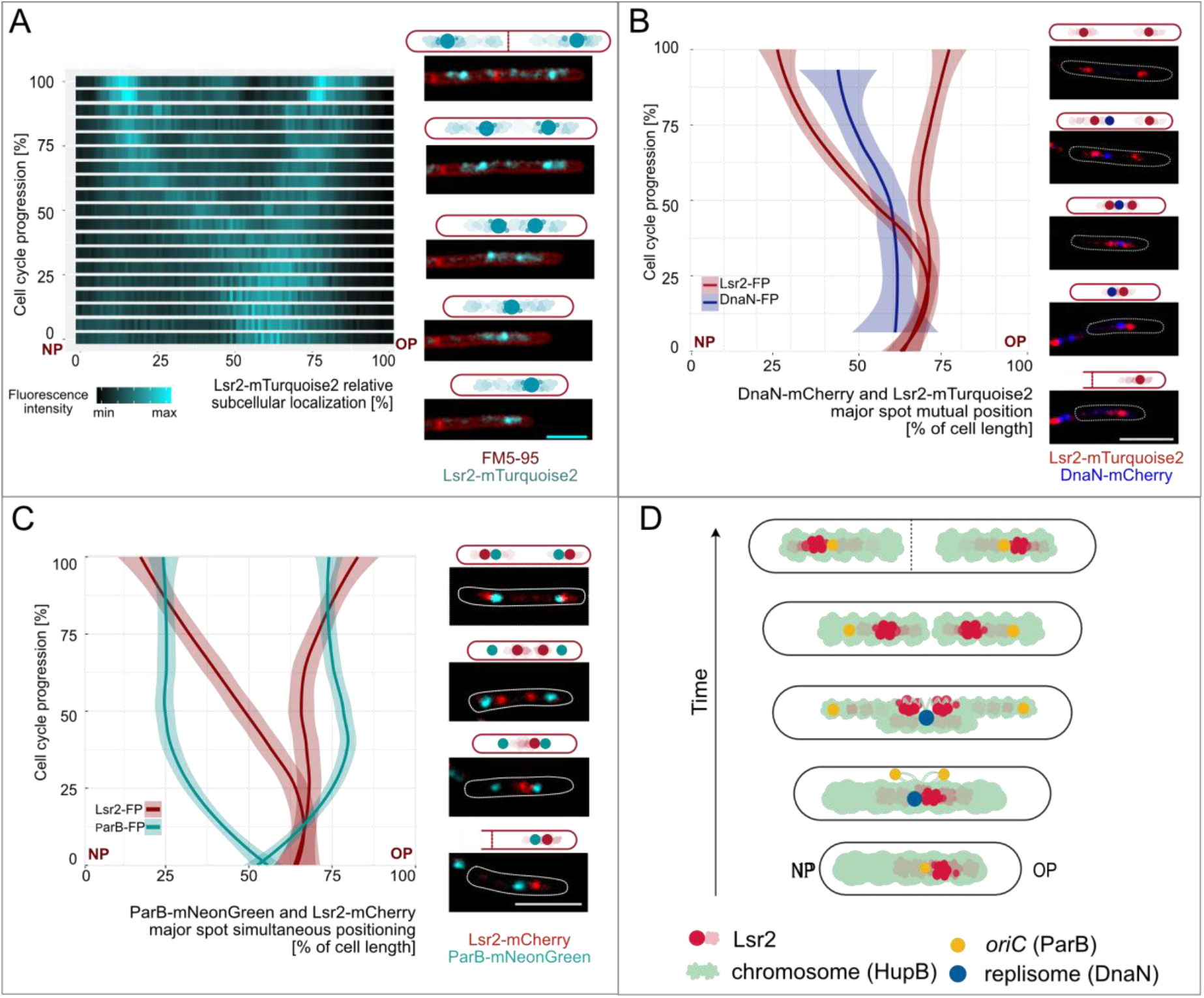
Subcellular localization of Lsr2-FP during the mycobacterial cell cycle. **(A)** Kymograph representing real-time Lsr2-mTurquoise2 localization (n = 5, left panel) and micrographs of representative FM5-95-stained Lsr2-mTurquoise2 cells. Bar, 2 μm (right panel). **(B and C)** Simultaneous positioning of the Lsr2-FP major foci and DnaN-mCherry (replisome) or ParB-mNeonGreen (*ori*C/segrosome) signals over time (left panels). Also shown are graphs representing the averaged position of fluorescent proteins (dark-colored line) and the standard deviation (light-colored ribbon, n = 5). For most of the C-period, in contrast, the major Lsr2-FP focus (foci) accompanies (accompany) the replisomes. Lsr2-FP and the segrosome(s) localized near one another at the beginning and end of the cell cycle. The images on the right of the sections show time-lapse microscopic images of representative Lsr2-mTurquoise2_DnaN-mCherry or Lsr2-mCherry_ParB-mNeonGreen cells. Bar, 5 μm. NP – new pole, OP – old pole. **(D)** Cartoon illustrating the subcellular localization of Lsr2 during the mycobacterial cell cycle.

To localize Lsr2 along the chromosome, we constructed a strain producing HupB-EGFP (as a chromosome marker) and Lsr2-mCherry fusion proteins from their native chromosomal loci (for details see Table S1). TLFM analysis of the HupB-EGFP_Lsr2-mCherry cells showed that during the entire cell cycle, the major Lsr2 focus (or two foci) and the weaker diffused foci were localized within the area occupied by the nucleoid (Fig. S3, Video S1). The major focus was duplicated at approximately the 1/3 point of cell cycle progression, and thereafter both of the duplicated foci moved towards opposite cell poles (Fig. 6A). Interestingly, the focus closest to the old cell pole was positioned at a relatively constant distance from the edge of the chromosome (20 +/−10% of the chromosome length, n = 390) at all-time points, while the second one traversed the chromosome to reach the opposite edge (Fig. S3).

Since the subcellular localization of the two major Lsr2-mCherry foci during the cell cycle resemble the positioning of segrosomes (i.e., ParB-FP complexes (Ginda et al., 2017)) we decided to track both proteins in the ParB-mNeonGreen_Lsr2-mCherry strain (for details see Table S1) and analyze their dynamics at different time intervals (Fig. 6C). TLFM observations revealed that at the beginning of the cell cycle, the Lsr2-mCherry and ParB-mNeonGreen foci were positioned slightly off-center, closer to the old cell pole, and they were relatively close together. Immediately following the initiation of replication, the ParB-mNeonGreen complex was duplicated and both foci were rapidly segregated towards the opposite cell poles; in contrast, the Lsr2-mCherry focus remained around the midcell for 53 +/− 9 min (n = 50) after the ParB-mNeonGreen duplication. This was followed by duplication of the major Lsr2-mCherry focus and the subsequent movement of these foci to the opposite cell poles. Shortly before the division of the cell, the translocated Lsr2-mCherry foci exhibited polar localizations that were more pronounced than those of the ParB foci (Fig. 6C, Video S1). Thus, fluorescent foci derived from Lsr2-mCherry and ParB-mNeonGreen were localized close together only at the beginning and end of the cell cycle, when chromosome replication did not occur (Fig. 6C,D).

Because the lack of Lsr2 protein altered the C-period duration and replisome dynamics, we decided to explore the localization of Lsr2-FP in the background of chromosome replication. For this purpose, we constructed a strain that simultaneously produced Lsr2-mTurquoise2 and DnaN-mCherry (for details see Table S1). Previous reports showed that *M. smegmatis* replisomes localize near the midcell and are placed asymmetrically on the chromosome (Hołówka et al., 2018; Trojanowski et al., 2015). At the time of replication initiation, the replisomes are assembled close to the edge of the chromosome in the old-pole-proximal cell half; before the termination of replication, the replisomes migrate toward the new-pole-proximal cell half. TLFM analysis of the Lsr2-mTurquoise2_DnaN-mCherry strain showed that both proteins localized close to one another for approximately the first half of the C-period. After the duplication of Lsr2-mTurquoise2, replisomes were observed in the vicinity of the new-pole-proximal major focus of Lsr2-mTurquoise (Fig. 6B, Video S1).

Together, these findings reveal that the major Lsr2-FP and ParB-FP complexes are localized relatively close together at the beginning and end of the cell cycle, and the main Lsr2-FP foci accompany the replisomes for most of the C-period.

### The oligomerization domain critically influences the subcellular localization of Lsr2

Previously, ChIP-on-chip analysis (B. R. G. Gordon et al., 2010) showed that Lsr2 binds to roughly a few hundred sites that are evenly distributed along the *M. smegmatis* chromosome. Since Lsr2-FP was visualized mainly as one or two discrete bright foci *in vivo*, we expected that Lsr2 might use its N-terminal oligomerization domain to bridge/link distal sites on the chromosome and thereby form higher-order nucleoprotein complexes. To test this hypothesis, we constructed strains producing either Lsr2-Dendra2 or its truncated form lacking the N-terminal oligomerization domain (Lsr2ΔNTD-Dendra2) (for details see Table S1, and Fig. 7B) from the native chromosomal locus. The phenotype of the Lsr2ΔNTD-Dendra2 strain resembled that of the Δ*lsr2* strain: it formed round and smooth colonies on 7H10 or NB agar plates and was not able to form biofilm (Fig. S1).

**Fig. 7.**
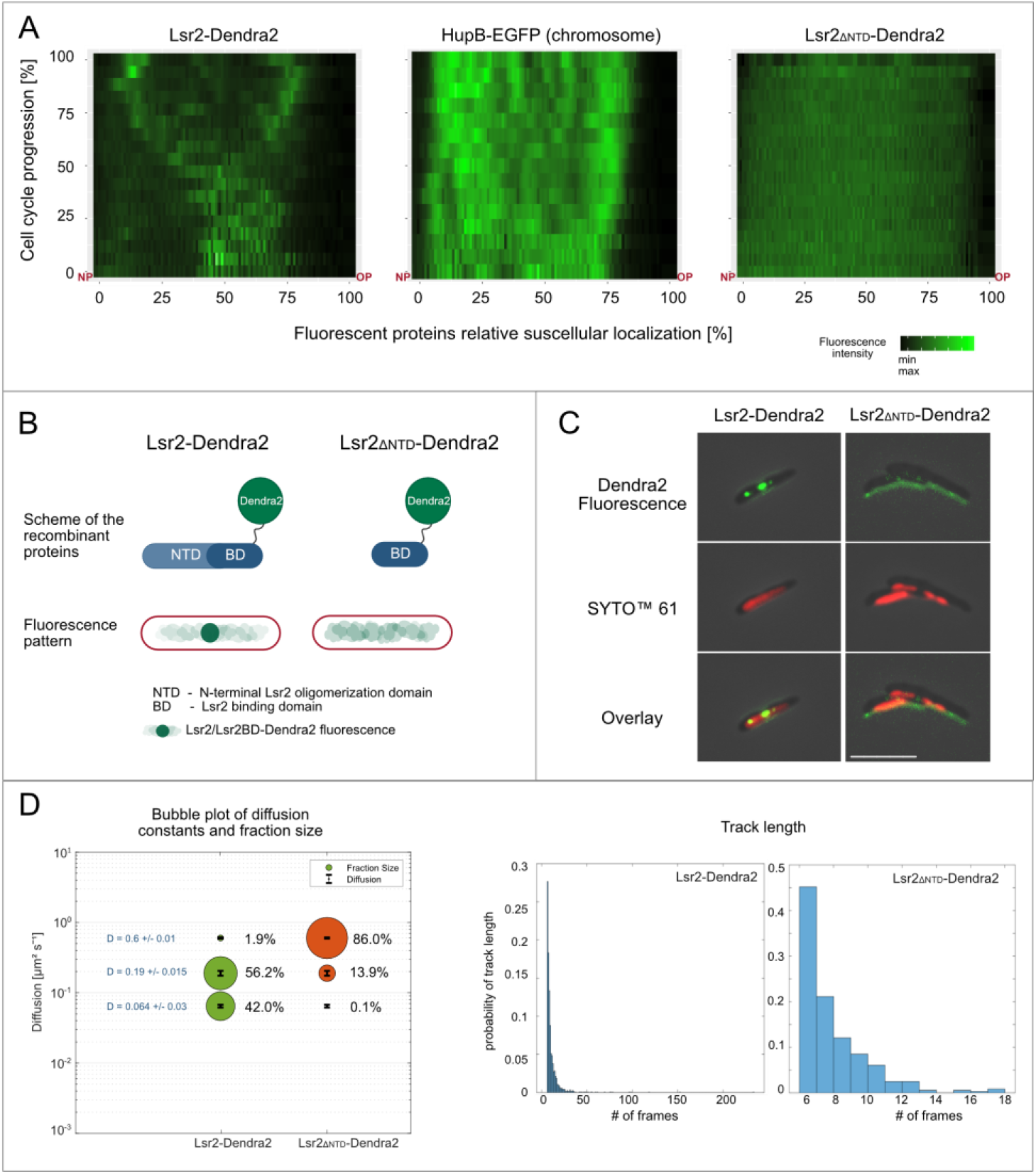
The influence of the N-terminal oligomerization domain on the subcellular localization and DNA binding of Lsr2. **(A)** Fluorescence profiles of Lsr2ΔNDT-Dendra2, HupB-EGFP (chromosome), and Lsr2-Dendra2 during the cell cycle (n = 5). The truncated version of Lsr2 is observed as diffused fluorescence along the cell. NP – new pole, OP – old pole. **(B)** Graphic representation of full-length and truncated Lsr2 fused with Dendra2 (top panel). Fluorescence patterns of both proteins in *M. smegmatis* cells (bottom panel). **(C)** Microscopic images of representative Lsr2-Dendra2 and Lsr2ΔNDT-Dendra2 cells stained with the nucleic acid dye, SYTO^TM^ 61. Bar, 5 μm. **(D)** PALM analysis of the mobilities of Lsr2-Dendra2 and Lsr2ΔNDT-Dendra2 proteins. The bubble plot comparison of diffusion constants and fraction sizes shows that the ratio of diffusing versus immobile particles was higher for the truncated form than for the full-length protein (left panel). The histograms represent the track length of full-length and truncated Lsr2-Dendra2.

Microscopic analysis revealed that, unlike Lsr2-Dendra2, Lsr2ΔNTD-Dendra2 was visible as a diffused fluorescence signal spread along the cell (Fig. 7A,C). We did not observe any individual Lsr2ΔNTD-Dendra2 focus at any point during the cell cycle. This suggests that the N-terminal domain of Lsr2 is indispensable for the formation of nucleoprotein complexes *in vivo.* Moreover, since a substantial fraction of Lsr2ΔNTD-Dendra2 was presumably not associated with the chromosome (see Fig. 7AC, compare HupB-EGFP and Lsr2ΔNTD-Dendra2), we postulate that the lack of the N-terminus may also reduce the DNA-binding affinity of the truncated Lsr2 protein *in vivo*. To verify our hypothesis, we used photoactivated localization microscopy (PALM) to visualize single molecules of Lsr2-Dendra2 and Lsr2ΔNTD-Dendra2 and analyze their mobility. Our results showed that the particles of both proteins were heterogeneous in their mobility. For full-length Lsr2-Dendra2 protein, we observed mainly two subpopulations: static molecules (diffusion constant, D = 0.064 +/− 0.003 μm^2^s^−1^) and slow-mobility molecules (D = 0.190 +/− 0.015 μm^2^s^−1^), which represented 42% and 56% of the population, respectively (2060 tracks from 91 cells were analyzed). Only a marginal fraction of the tracks (<2%) showed high mobility (D = 0.60 +/− 0.01 μm^2^s^−1^). In contrast, 86% (n = 365 tracks obtained from 92 cells) of the truncated protein molecules (Lsr2ΔNTD-Dendra2) exhibited high mobility, and only approximately 14% exhibited slow mobility (with D values corresponding to that of full-length Lsr2-Dendra2, Fig. 7D). Notably, we collected significantly fewer tracks for the truncated Lsr2, mainly because there were fewer points per track compared to those of the full-length protein (only tracks consisting of at least five points were included in the analysis). We observed the same tendency regardless of the exposure time applied during data acquisition. Analysis of dwell times (the period during which a particle resides inside a defined radius, which in our case corresponded to the time when Lsr2-Dendra2 or its truncated form remained bound to the chromosome) showed that all molecules of a given protein (fast vs. slow diffusing) exhibited the same dwell times. However, the dwell times calculated for Lsr2ΔNTD-Dendra2 were 2-fold lower than those obtained for full-length Lsr2-Dendra2 (47 +/− 4 ms vs. 82 +/− 2 ms, respectively).

In sum, our TLFM and PALM analyses demonstrated that the N-terminal domain of Lsr2 is essential for the ability of Lsr2 to form large nucleoprotein complexes *in vivo*.

## Discussion

The Lsr2 protein plays both architectural and regulatory roles and is highly conserved throughout the Actinobacteria, including the pathogenic mycobacteria and the antibiotic-producing *Streptomyces*. Lsr2 is a pleiotropic regulator of gene expression that acts mainly as a repressor. In *M. tuberculosis,* it regulates genes that are involved in multiple cellular processes, such as cell wall synthesis, antibiotic resistance, and virulence (Colangeli et al., 2009; Leistikow et al., 2010), while in *S. venezuelae*, it represses genes that are involved in the synthesis of cryptic specialized metabolites (Gehrke et al., 2019). However, there was no previously published information regarding the impact of Lsr2 on the cell cycle or cell morphology of mycobacteria or *Streptomyces*. Here, we demonstrate that in mycobacteria, Lsr2 has a profound effect on cell morphology and cell envelope properties. Moreover, real-time visualization of Lsr2-FP in single cells revealed that it forms a nucleoprotein complex that exhibits a highly dynamic localization during the cell cycle.

### Lsr2 influences single-cell morphology and cell envelope properties

Consistent with previous analyses showing that *M. smegmatis* strains lacking the *lsr2* gene demonstrated profound changes in colony morphology (Arora et al., 2008; Chen et al., 2006; Gordon et al., 2008), we observed that the Δ*lsr2* strain formed round, smooth colonies, was unable to form biofilm and spread on the surface of solid medium (Fig. S1). Our single-cell AFM-based studies showed that Δ*lsr2* cells have smoother surfaces than WT cells (Fig. 2B, bottom panel). We expect that this is directly related to the changes in the cell envelope composition (mainly its outer membrane). Indeed, microarray analysis (Colangeli et al., 2007) confirmed by the qRT-PCR (Fig. 4A) revealed that the expression of MSMEG_4727 gene involved in the lipooligosaccharides (LOSs) synthesis is significantly higher in *M. smegmatis* strain lacking the Lsr2 protein. Moreover, TLC and MALDI-TOF MS analyses of crude lipids confirmed that the Δ*lsr2* strain, in contrast to the wild type, produces LOSs (Fig. S2C). LOSs contain a hydrophilic carbohydrate moiety and are found in the outer membrane of some mycobacterial species (Jankute et al., 2017). In *M. kansasii,* lipooligosaccharides have been also shown to be associated with colony smoothness, sliding motility, and inability to form a biofilm. Interestingly, unlike smooth variant of *M. kansasii*, the rough, LOSs-deficient variants of *M. kansasii* are capable of causing a chronic systemic infection in mice (Collins & Cunningham, 1981; Belisle & Brennan, 1989). Notably, recent studies suggested that *M. tuberculosis* during the evolution became more hydrophobic by deletion of genes involved in the synthesis of polar lipids including LOSs. It is believed that such a change presumably enhances the capability for aerosol transmission, affecting virulence and pathogenicity (Johansen, Herrmann, & Kremer, 2020).

Moreover, the Δ*lsr2 M. smegmatis* strain exhibits higher susceptibility to vancomycin (Fig.3). This phenomenon might reflect the increased antibiotic penetration through the cell envelope. This explanation seems probable since the highly hydrophilic carbohydrate part of LOSs might promote penetration of hydrophilic antibiotic, such as vancomycin.

Our single-cell microscopic observations revealed that mycobacterial cells lacking the Lsr2 protein exhibit morphological abnormalities, including shortening and widening of the cell (Fig. 1). We speculate that the shortening of the *Δlsr2* cell length is solely due to their reduced cell elongation rate (Fig. 5A). Consistent with this hypothesis, we found that despite the difference in the elongation rate between the Δ*lsr2* and WT strains, their doubling times (B+C+D) were similar (Fig. 5B). We also expect that the shortening of cell length and the changes in cell envelope composition/properties have other consequences. For example, macromolecular crowding is likely to be increased within the shorter cells (Mourão, Hakim, & Schnell, 2014). Our AFM data supported this assumption: Young’s modulus, which reflects the turgor pressure (related to cell stiffness) was higher for Δ*lsr2* versus WT cells (Fig. 2, top panel). A high turgor pressure is frequently associated with increase in cytoplasmic concentration, macromolecular crowding, and depletion forces (Arnoldi et al., 2000; Rojas & Huang, 2018).

### Lsr2 forms large dynamic nucleoprotein complexes that are presumably involved in the organization of DNA strands during replication and segregation

While the biochemistry of the Lsr2 protein has been well studied (Chen et al., 2008; Gordon et al., 2011; Qu et al., 2013; Summers et al., 2012), its subcellular localization and influence on the mycobacterial cell cycle were previously unknown. Our real-time imaging analysis revealed that the fluorescently tagged Lsr2 (Fig. 6A) formed one or two bright major foci, along with several minor complexes and dispersed fluorescence that presumably reflect binding to the single cluster of Lsr2 target sequences and disassembly of nucleoprotein complexes during ongoing replication, respectively. Since Lsr2 was observed *in vivo* as a large focus/foci within the area occupied by the chromosome (Fig. S3, 7C), we assume that this protein forms nucleoprotein complexes in mycobacterial cells. Their localization is dynamic during the cell cycle. At the beginning of the cell cycle, the single bright Lsr2 focus is localized slightly asymmetric in relation to the midcell (closer to the old cell pole) near the *oriC* region (i.e., the ParB-FP complex). Soon after the initiation of DNA replication, the newly replicated *oriC* regions are rapidly segregated towards the opposite cell poles (Ginda et al., 2017; Hołówka et al., 2018; Santi & McKinney, 2015; Trojanowski et al., 2015), while Lsr2 remains localized in the vicinity of replisomes as a single bright focus. After approximately 1/3 of the cell cycle has been completed, the major Lsr2 focus splits into two foci, which then gradually move towards the opposite cell poles where the segrosomes are already located. Thus, at the beginning and end of the cell cycle, the Lsr2 foci are localized at cell poles near the segrosomes (Fig. 6C,D, Video S1). For most of the C-period, in contrast, the Lsr2 foci occupy midcell positions near the replisomes. The main Lsr2 focus splits into two foci when approximately half of the chromosome has been copied (see Fig. 6B). Interestingly, our results revealed that the two major Lsr2 foci move asymmetrically - with one traveling a greater distance (this one moving to the new pole). This asymmetric translocation mode of the Lsr2 foci presumably reflects asymmetric action of the segregation machinery (Ginda et al., 2017; Holówka et al., 2018) and the unequal division and growth of mycobacterial cells.

Since the major focus (or foci) of Lsr2 is (are) localized near the replisomes (Fig. 6B), we speculate that this protein influences chromosome replication. Indeed, the duration of DNA replication is shorter in Δ*lsr2* cells (Fig. 5B), presumably due to the lack of spatial obstacles (i.e., Lsr2 nucleoprotein complexes). In the WT strain, such obstacles might need to be removed from the DNA ahead of the replication machinery. The presence of Lsr2 nucleoprotein complexes in the vicinity of replisomes may also suggest that, similar to HN-S in *E. coli* (Helgesen, Fossum-Raunehaug, & Skarstad, 2016), the Lsr2 protein is involved in organizing the sister DNA stretches that are moved away from the replication machinery. Moreover, at around the halfway point of chromosome replication, the DNA must undergo significant conformational changes due to reorganization of the Lsr2 nucleoprotein complex (the Lsr2 focus splits into two foci). We speculate that the Lsr2-mediated dynamic reorganization of DNA both behind and ahead of the moving replication fork may affect the dynamics of replisomes. In fact, we observed that the replisomes in WT cells frequently divided and merged, and that this was in contrast to the sparse splitting of replisomes in the Δ*lsr2* cells (Fig. 5C, D). Meanwhile, the Δ*lsr2* cells might have difficulty in organizing sibling chromosomes; the absence of Lsr2 presumably causes local chromosome disorganization, which could extend the B+D periods and delay chromosome initiation in the daughter cells (Fig. 5B).

Given that the localization of the major Lsr2 focus on the chromosome is highly dynamic (Fig. S3, Video S1), we speculate that Lsr2 nucleoprotein complexes need to be dynamically assembled and disassembled during the course of chromosome replication, to organize the newly replicated DNA and to enable efficient progression of the DNA replication machinery, respectively. The Lsr2 protein presumably bridges different regions of chromosomes depending on the progress of DNA replication. Notably, Chip-ChIP found that the few hundred Lsr2 binding sites are evenly distributed along the *M. smegmatis* chromosome (Gordon et al., 2010). However, the analysis was carried out on non-synchronized cells.

Previous *in vitro* studies showed that the N-terminal domain of Lsr2 is involved in oligomerization (Chen et al., 2008; Summers et al., 2012), enabling Lsr2 to form large nucleoprotein complexes by bridging distant DNA fragments. To elucidate the *in vivo* role of the N-terminal domain (NTD) of Lsr2, we constructed fluorescent reporter strains that produced a truncated form of Lsr2 lacking the NTD, Lsr2ΔNTD-Dendra2, or the wild-type protein, Lsr2-Dendra2 (Fig. 7B), and compared the subcellular localizations of the fusion proteins using TLFM and high-resolution PALM (photoactivated localization microscopy). Previous *in vitro* experiments had demonstrated that Lsr2ΔNTD binds DNA (Gordon et al., 2011).

Our TLFM analysis showed that Lsr2ΔNTD-Dendra2 was visible as diffuse fluorescence along the cell (Fig. 7A,C), whereas Lsr2-Dendra2 exhibited the foci described above. Notably, we did not observe any individual Lsr2ΔNTD-Dendra2 focus at any point during the cell cycle. In addition, the PALM analysis revealed that the ratio of diffusing versus immobile particles was considerably higher for the truncated form than for the wild-type protein (Fig. 7D), suggesting that a substantial fraction of the Lsr2ΔNTD-Dendra2 particles may be present in the cytoplasm rather than bound to the nucleoid. We note, however, that the high mobility of the Lsr2ΔNTD-Dendra2 particles may be the result of unstable (and transient) DNA binding and competition with other DNA binders, including NAPs. Thus, our *in vivo* observations are consistent with the previous *in vitro* experiments and support the hypothesis that the N-terminus of the Lsr2 protein is indispensable for the formation of the massive nucleoprotein complexes that are visible as bright fluorescent foci *in vivo*.

In sum, we herein used a suite of complementary microscopic techniques (i.e., TLFM, AFM, PALM) to elucidate the influence of Lsr2 on single-cell morphology and the cell cycle of *M. smegmatis,* which is a model organism for pathogenic mycobacteria including *M. tuberculosis*. Our results indicate that Lsr2 exerts profound effects on cell morphology, cell envelope properties, and the dynamics of chromosome replication. *In vivo*, Lsr2 forms a nucleoprotein complex that exhibits a highly dynamic localization during the cell cycle. Moreover, the Lsr2 N-terminal domain, which is involved in oligomerization, is indispensable for the *in vivo* formation of nucleoprotein complexes. Given that the Lsr2 protein is essential during the infection of *M. tuberculosis* (Colangeli et al., 2007; McAdam et al., 2002; Sassetti, Boyd, & Rubin, 2003) and exhibits pleiotropic activities, Lsr2 appears to be an attractive target for the development of new antimicrobials.

## Materials and Methods

### DNA manipulations, bacterial strains, and culture conditions

All plasmids used for transformation of *M. smegmatis* mc^2^ 155 were propagated in the *E. coli* DH5α strain. *E. coli* was grown in LB broth or on LB agar plates (Difco) supplemented with the proper antibiotic(s) and/or other compounds (100 μg/ml ampicillin, 50 μg/ml kanamycin, 50 μg/ml hygromycin, 0.004% X-Gal [5-bromo-4-chloro-3-indolyl-α-D-galactopyranoside]), accordingly to standard procedures (Sambrook, Fritsch, & Maniatis, 1989). *M. smegmatis* strains were grown in 7H9 broth supplemented with 10% OADC (oleic acid-albumin-dextrose-catalase; BD) and 0.05% Tween 80, or on 7H10 agar plates (Difco) supplemented with 10% OADC, 0.5% glycerol, 0.004% X-Gal, 2% sucrose and/or kanamycin, and 50 μg/ml hygromycin. DNA manipulations were carried out using standard protocols (Sambrook et al., 1989). Reagents and enzymes were obtained from Thermo Fisher, Roth, and Merck (Sigma-Aldrich). Oligonucleotides were synthesized by Genomed (Poland) or Merck (Sigma-Aldrich), and sequencing was performed by Genomed or Microsynth (Germany). To determine the growth curve for *M. smegmatis* strains in optimal conditions and during exposure to vancomycin, cells were grown at 37°C in a final volume of 300 μl 7H9 (supplemented OADC and Tween80), and optical density measurements were taken at 10-min intervals for 24–72 hours using a Bioscreen C instrument (Growth Curves US). Bacterial strains, plasmids, and oligonucleotides are listed in Table S1-3 in the supplemental material. The construction of the *M. smegmatis* mc^2^ 155 mutant strains is detailed in Supplemental Experimental Procedures.

### Microscopy and cell staining

Snapshot imaging of *M. smegmatis* was performed using cells cultured to log phase (optical density at 600 nm [OD600] ~ 0.6). Cells were directly plated on an agar pad containing FM5-95 (0.5 μg/L; Thermo Fisher) for membrane staining or incubated with SYTO^TM^61 (1.6 μM) for 15 minutes prior to being plated on the agar pad for nucleoid acid staining. Real-time analyses were performed on solid medium in ibidi micro dishes and in liquid medium using a CellASIC ONIX platform and compatible B04A plates (Merck), as described previously (Hołówka et al., 2017; Trojanowski et al., 2015; Trojanowski et al., 2019). In both cases, early log phase (OD600 ~ 0.2 to 0.4) *M. smegmatis* cultures grown in liquid medium were used. For membrane staining, cells were plated on an ibidi μ-dish (35 mm, low) with solid medium (7H10 supplemented with OADC) containing FM5-95 (0.5 μg/L). For NADA (fluorescent D-amino acid; Torcis Bio-Techne)/TMR-Tre (fluorescent trehalose; Torcis Bio-Techne) staining (Kuru et al., 2012; Kuru et al., 2015), cells loaded into the observation chamber were cultured in 7H9 (supplemented with OADC and Tween80). They were then subjected to 2-minute pulse flows of 7H9-OADC-Tween 80 medium supplemented with NADA (0.5 mM) or TMR-Tre (100 μM) every hour. For Vancomycin-BODIPY (BODIPY^TM^ FL Vancomycin; Thermo Fisher) staining, cells loaded into the observation chamber were exposed to fresh 7H9-OADC-Tween 80 for 5 hours, followed by 7H9-OADC-Tween 80 supplemented with 0.5 μg/ml Vancomycin-BODIPY for 5 hours, and then 7H9-OADC-Tween 80 supplemented with 1 μg/ml Vancomycin-BODIPY for 5 hours (Fig. 3). All microfluidic experiments were performed under constant pressure (1.5 psi). Images were recorded at 10 minutes intervals using a Delta Vision Elite inverted microscope equipped with a 100× oil immersion objective and an environmental chamber set to 37°C.

Pictures were analyzed using the Fiji and R software packages (R Foundation for Statistical Computing, Austria; http://www.r-project.org), including the ggplot2 package (Wickham, 2009). For all measurements, a two-sided parametric Student’s t-test was applied. To avoid the generation of false assumptions in the case of nonnormal distributions, the statistical significance of differences in the measured values was confirmed with the nonparametric two-sided Wilcoxon test with minimum confidence intervals of 0.995.

### Atomic force microscopy (AFM)

To estimate the roughness profile, *M. smegmatis* cultures were grown in 7H9-OADC-Tween 80 medium, centrifuged (6000 rpm for 5 minutes), washed with PBS and water, smeared onto microscopic slides, and dried. AFM measurements were conducted using the PeakForce Tapping mode (scan rate, 1.5 Hz; amplitude, 100 nm) on a BioScope Resolve AFM microscope (Bruker). ScanAsyst-Fluid+ cantilevers (Bruker) were used for AFM scanning (resonant frequency, 150 kHz; spring constant, 0.7 N/m). The Ra parameter (arithmetic mean deviation of the roughness profile) was calculated from the cell surface area using the NanoScope Analysis 1.9 software (Bruker).

To estimate Young’s modulus, *M. smegmatis* cultures were grown in 7H9-OADC-Tween 80 medium. Two milliliters of bacterial culture were centrifuged (6000 rpm for 5 minutes), washed twice with 7H9-OADC (to remove the Tween 80) and resuspended in 50 μl 7H9-OADC. Microscope slides were prepared as previously described (Eskandarian et al., 2017) by mixing polydimethylsiloxane (PDMS; Sylgard 184; Dow Corning) at a ratio of 15:1 (elastomer:curing agent). Air bubbles were removed from the mixture under negative pressure for 20 minutes. The PDMS mixture was dropped onto the microscope slides (VWR), which were spin-coated and baked at 80°C for 10 minutes before use. A concentrated bacterial pellet was deposited on the surface of a PDMS-coated slide and incubated for 10 minutes. Then, the PDMS-coated slide was washed with 7H9-OADC to remove non-immobilized cells. AFM measurements were conducted in 7H9-OADC medium. The Young’s modulus was estimated using the PeakForce QNM scanning mode (scan rate, 1 Hz; amplitude, 100 nm) with PFQNM-LC-A-CAL cantilevers (resonant frequency, 45 kHz; spring constant, 0.1 N/m; Bruker). Young’s modulus was measured at five points of each cell using the NanoScope Analysis 1.9 software, and then the mean modulus value was calculated for *M. smegmatis* WT and Δ*lsr2* cells.

### Single-particle tracking

Lsr2-Dendra2 and Lsr2ΔNTD-Dendra2 cells were cultured to mid-log phase in rich medium (7H9 supplemented with 10% ADC and 0.05% Tween 80). Immediately before imaging, cells were spread onto agar pads (1% agarose in 7H9 poured into 1.0×1.0-cm GeneFrames; Thermo Fisher Scientific) and covered with a clean 22×22 0.17-mm coverslip. Imaging was carried out using a Zeiss Elyra P.1 inverted microscope equipped with an Andor iXon 897 back-thinned EMCCD camera and a 100x/1.46 Oil DIC objective. The samples were prebleached and the images were recorded using a 10.3-ms exposure per frame (561-nm laser, 30% intensity, 10k frames in total) and increasing 405-nm excitation (continuous increase from 0.002 to 1%). Data were analyzed using Fiji, Oufti, and SMtracker software packages.

## Acknowledgments

The work was financed by a grant (OPUS 2017/25/B/NZ1/00657) from the National Science Center (Poland). We are grateful to Marie Elliot for her critical reading of the manuscript.

